# Identifying barriers to gene flow and hierarchical conservation units from seascape genomics: a modelling framework applied to a marine predator

**DOI:** 10.1101/2021.10.25.465682

**Authors:** Germain Boussarie, Paolo Momigliano, William D. Robbins, Lucas Bonnin, Jean-François Cornu, Cécile Fauvelot, Jeremy J. Kiszka, Stéphanie Manel, David Mouillot, Laurent Vigliola

**Affiliations:** UMR ENTROPIE (IRD, UR, UNC, IFREMER, CNRS), Centre IRD de Nouméa, Nouméa, Nouvelle-Calédonie, France; UMR MARBEC (CNRS, IRD, IFREMER, UM), Université de Montpellier, Montpellier, France; Organismal and Evolutionary Biology Research Programme, University of Helsinki, Helsinki, Finland; Wildlife Marine, Perth, Western Australia, Australia; Department of Environment and Agriculture, Curtin University, Perth, Western Australia, Australia; School of Life Sciences, University of Technology Sydney, Sydney, New South Wales, Australia; Department of Biodiversity, Conservation and Attractions, Marine Science Program, Biodiversity and Conservation Science, Kensington, Western Australia, Australia; UMR ENTROPIE (IRD, UR, UNC, IFREMER, CNRS), Laboratoire d’Océanographie de Villefranche, Villefranche-sur-Mer, France; UMR LOV (Sorbonne Université, CNRS), Villefranche-sur-Mer, France; Institute of Environment, Department of Biological Sciences, Florida International University, Miami, Florida, USA; CEFE (Univ Montpellier, CNRS, EPHE-PSL University, IRD), Montpellier, France

**Author notes:** **Correspondence** Germain Boussarie, Institut de Recherche pour le Développement (IRD), Centre IRD de Nouméa, BP A5, 98848 Nouméa cedex, New Caledonia. These authors contributed equally to this work (David Mouillot, Laurent Vigliola). **Ethics** Genetic samples in New Caledonia were collected under permits to Laurent Vigliola by the Government of New Caledonia (permit #2015-1351/GNC), the Northern Province of New Caledonia (permit #60912-1508-2015/JJC) and the Southern Province of New Caledonia (permits #479-2016/ARR/DENV and #2093-2016/ARR/DENV). Great Barrier Reef Marine Park permits include G01/356, G02/021 and G03/8048.1. **Author contributions** GB, LV, DM, PM designed the study, GB, LV, LB, WR, JK, PM collected the data, GB analyzed the data and wrote the first draft with inputs from LV, DM, PM, all authors contributed to writing, review and editing.

**Keywords:** Seascape genomics, conservation, gene flow, circuit theory, isolation-by-resistance, reef sharks

## Abstract

The ongoing decline of large marine vertebrates must be urgently mitigated, particularly under increasing levels of climate change and other anthropogenic pressures. However, characterizing the connectivity among populations remains one of the greatest challenges for the effective conservation of an increasing number of endangered species. Achieving conservation targets requires an understanding of which seascape features influence dispersal and subsequent genetic structure. This is particularly challenging for adult-disperser species, and when distribution-wide sampling is difficult. Here, we developed a two-step modelling framework to investigate how seascape features drive the genetic connectivity of marine species without larval dispersal, to better guide the design of marine protected area networks and corridors. We applied this framework to the endangered grey reef shark, *Carcharhinus amblyrhynchos*, a reef-associated shark distributed across the tropical Indo-Pacific. In the first step, we developed a seascape genomic approach based on isolation-by-resistance models involving circuit theory applied to 515 shark samples, genotyped for 4,491 nuclear single-nucleotide polymorphisms, to explore which parameters drive their population genetic differentiation. We show that deep oceanic areas act as strong barriers to dispersal, while proximity to habitat facilitates dispersal. In the second step, we predicted the resulting genetic differentiation across the entire distribution range of the species, providing both local and global-scale conservation units for future management guidance. We found that grey reef shark populations are more fragmented than expected for such a mobile species, raising concerns about the resilience of isolated populations under high anthropogenic pressures. We recommend the use of this framework to identify barriers to gene flow and to help in the delineation of conservation units at different scales, together with its integration across multiple species when considering marine spatial planning.

## 1 INTRODUCTION

Marine ecosystems across the globe are under increasing pressure due to habitat fragmentation, overexploitation, and climate change (Albouy et al., 2020; McCauley et al., 2015; Young et al., 2016). Due to their conservative life-history traits of low reproductive rates, high longevity, and slow growth, large marine vertebrates such as marine mammals, seabirds, and elasmobranchs are particularly vulnerable to human-induced mortalities: their rate of extinction is indeed higher due to fisheries (including bycatch), habitat disturbance, and pollution (Estes et al., 2016; MacNeil et al., 2020; McClenachan et al., 2016; Yan et al., 2021). Their effective protection is an unprecedented challenge that must be addressed in the coming decades (Duarte et al., 2020; Sala et al., 2021).

The implementation of effective conservation measures for large marine vertebrates requires that space use by these potentially highly mobile species is taken into account (Harrison et al., 2018; Jacoby et al., 2020), and to better understand the factors driving connectivity among populations in increasingly fragmented seascapes (Balbar and Metaxas, 2019; McRae and Beier, 2007). Indeed, through the exchange of genes, connectivity plays a vital role in maintaining thriving natural populations (Cowen and Sponaugle, 2009; Dunn et al., 2019; Jangjoo et al., 2016), ensuring biodiversity conservation and fisheries sustainability (Álvarez-Noriega et al., 2020; Edgar et al., 2014; Gaines et al., 2010; Krueck et al., 2017).

Population connectivity of most marine animals depends on a dispersive planktonic larval phase. This life-history stage can last from days to months, and larval dispersal can be modelled using biophysical or genetic frameworks (Bryan-Brown et al., 2017; Harrison et al., 2020; Manel et al., 2019). Genetic connectivity, a measure of the degree to which gene flow affects evolutionary processes among populations, has been widely studied among larval dispersers (*e*.*g*. Benestan et al., 2021), since gene flow plays a key role in maintaining genetic diversity and healthy populations able to adapt to a changing environment (Goetze et al., 2021; Lowe and Allendorf, 2010; Slatkin, 1987; Song et al., 2013).

In contrast, investigating the population or genetic connectivity of species whose dispersal is realized by adults is more challenging as adult connectivity cannot be modeled using the same oceanographic models (*e*.*g*. Boissin et al., 2019; Pazmiño et al., 2017; Pirog et al., 2019). Yet, the question of which factors drive nektonic adult connectivity has received less attention, while its knowledge is just as important for those species relying on dispersal of larger individuals to maintain connectivity (Momigliano et al., 2015). Indeed, little is known about which habitat, environmental and biogeographic features drive the connectivity among populations for adult dispersers across generations. Investigating these factors can provide key information to properly design corridors and networks of marine protected areas (MPA) (Almany et al., 2009; Balbar and Metaxas, 2019; Jacoby et al., 2020; Magris et al., 2014, 2018).

One approach to investigate the factors shaping genetic connectivity among populations and identify subsequent barriers to gene flow is the use of isolation-by-resistance (IBR) models (McRae, 2006). These models, popular in terrestrial ecology (Dickson et al., 2019), remain largely overlooked in the marine realm (Selkoe et al., 2016). Unlike isolation-by-distance (IBD) models, IBR models incorporate the effects of heterogeneous habitats on gene flow, thus they can account for the effect of seascape features on the genetic differentiation among populations and also make predictions for sites that have not been sampled (McRae and Beier, 2007). A combination of large empirical genetic datasets and modelled genetic differentiation could therefore be used to delineate conservation units (groupings of a species which contain sufficient biodiversity for persistence through subsequent generations) throughout the entire range of a species.

Separating a species’ range into conservation units can indeed identify key areas for dispersal along with populations under potential threats (Allendorf et al., 2010; Barbosa et al., 2018; Funk et al., 2012). Although the definition of conservation units has been debated (Lowe and Allendorf, 2010; Palsbøll et al., 2007; Waples and Gaggiotti, 2006), the hierarchical delineation of population subdivisions, based on genetic connectivity, can provide significant clues for both local and global management strategies (Barbosa et al., 2018; Dilts et al., 2016). Surprisingly, the degree of habitat fragmentation and the subsequent delineation of conservation units are poorly investigated in threatened and mobile marine species. Advances in genetic tools and computational power (Balkenhol et al., 2017; Barbosa et al., 2018; DiBattista et al., 2017; Funk et al., 2012; Schadt et al., 2010) now permit the development of models predicting how seascape features shape connectivity over a large scale at high spatial resolution (Leonard et al., 2017).

With no larval stage, the adult dispersion of sharks is of high importance (Hirschfeld et al., 2021), and shark conservation remains challenging with many species showing extensive geographic ranges spanning several countries (Dulvy et al., 2017, 2021; Pacoureau et al., 2021). Additionally, most shark species are highly vulnerable to fishing pressure given their life history traits, *e*.*g*. slow growth, late sexual maturity and low fecundity (Dulvy et al., 2014). Over one-third of chondrichthyans are threatened with extinction (*International Union for Conservation of Nature* Red List ; Dulvy et al., 2021), and some have undergone declines greater than 70% in abundance in the last few decades (Pacoureau et al., 2021; Roff et al., 2018). The size of MPAs is known to be a major driver of their protection effectiveness (Bonnin et al., 2021; Dwyer et al., 2020; Juhel et al., 2017, 2019), however these areas often only encompass a small proportion of each population’s distribution.

Here, we focused on the grey reef shark (*Carcharhinus amblyrhynchos*) as a model species to explore the potential offered by seascape genetics for the characterization and prediction of genetic connectivity of adult dispersers, the identification of barriers and resistance to dispersal, and possible implications for the spatial delineation of conservation units for management purposes. This species has strongly declined in non-protected reefs close to human habitation (Juhel et al., 2017, 2019; Robbins et al., 2006; Ruppert et al., 2017) and is now listed as *Endangered* on the IUCN Red List. It shows a high level of residency and small home range but adults can perform long-range movements (>700 km) along reefs and across oceanic waters (Bonnin et al., 2019, 2021; Espinoza et al., 2015a; White et al., 2017).

We followed a two-step approach to investigate the genetic connectivity of this near-threatened coral reef-associated predator and delineate hierarchical conservation units based on estimates of genetic connectivity. Firstly, we employed IBR modelling and electrical circuit theory (CT) (McRae, 2006; McRae et al., 2008) to determine how seascape features shape the genetic differentiation of this species, using an extensive genetic dataset of over 500 sharks collected in 17 locations across the Indian and Pacific oceans. We then used this modelling framework to delineate hierarchical conservation units across the whole species distribution range (Indo-Pacific), to better inform conservation strategies and identify the most vulnerable populations.

## 2 MATERIALS AND METHODS

### 2.1 Shark sampling and locations

We collected fin clips from 515 individual grey reef sharks across 17 locations in the Indian and Pacific Oceans (Figure 1). Samples from the Indian Ocean (n=99), Indonesia (n=24) and the Great Barrier Reef (n=48) were already described and genotyped in a published study (Momigliano et al., 2017). The remaining samples (n=344) were collected in the New Caledonian Archipelago, between June 2015 and November 2016, using barbless circle hooks for mouth-hooking and easy release after sampling. Sharks were caught on single lines to reduce bycatch and minimize handling and processing times.

**Figure 1.**
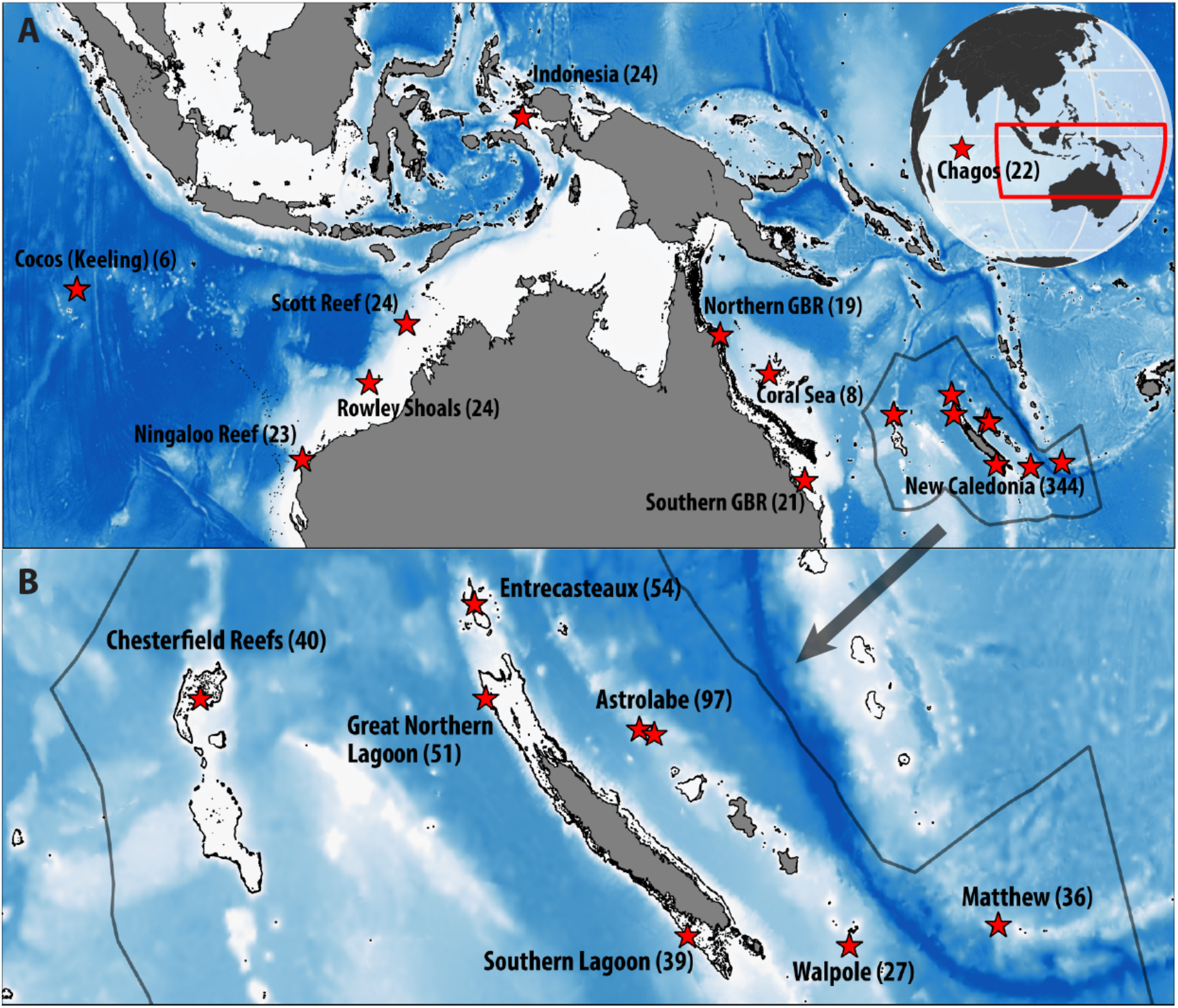
Maps of the 17 sampling locations where 515 grey reef shark samples were collected. (A) Global sampling locations (B) Detailed sampling at the scale of the New Caledonian archipelago (EEZ outlined in grey). The number of individuals sampled for SNP analysis from each location is in brackets.

### 2.2 Population genomics

We extracted DNA from fin clips using DNEasy Blood and Tissue kit (QIAGEN) for the 344 samples from New Caledonia. Each DNA solution was adjusted, after quality control, to 12-15 µL at 50 ng.µL^-1^ prior to DNA sequencing at Diversity Arrays Technology Pty. Ltd (Canberra, Australia), using DArTseq protocol (Sansaloni et al., 2011). Post-extraction laboratory protocols and SNP calling and filtering procedures used were the same as described in Momigliano et al. (2017), except for filtration to remove loci with minor allele frequencies, where we filtered SNPs for MAF>0.02 instead of 0.05, as sampling was extended to numerous additional locations.

After combining these SNPs with those of a previous study (Momigliano et al., 2017), outlier tests were used to filter loci for which genetic differentiation (*F*_ST_) is higher than expected under neutral processes only. We only included individuals from Australian, Indonesian, and New Caledonian sampling locations that showed little genetic differentiation at nuclear loci. We applied a combination of two methods to identify and exclude from further analyses these loci potentially under selection; OutFLANK (Whitlock and Lotterhos, 2015) and FLK, *i*.*e*. extensions of the Lewontin–Krakauer test that accounts for population co-ancestry (Bonhomme et al., 2010).

### 2.3 Population structure

We applied a Bayesian unsupervised clustering method (fastSTRUCTURE) to investigate genetic structure at neutral loci (Raj et al., 2014). fastSTRUCTURE implements an efficient algorithm for approximate inference of the admixture model from STRUCTURE (Pritchard et al., 2000). We ran fastSTRUCTURE with simple and logistic priors, at multiple numbers of clusters, K ranging from 1 to 10.

We also carried out Discriminant Analysis of Principal Components (DAPC) using the R package adegenet, with sampling location of each individual used as prior information (Jombart et al., 2010) to investigate patterns of genetic structure at neutral loci. The number of principal components (PCs) to retain for DAPC analyses was determined by cross-validation using a training set of 80% of the data and we therefore retained the number of PCs for which the obtained mean square error was the lowest.

### 2.4 Isolation-by-distance and isolation-by-resistance models

The relationship between genetic distance at neutral SNP loci (*F*_ST_) and geographic distance, *i*.*e*. isolation-by-distance (IBD) pattern, was investigated using multiple regression on distance matrices (MRM ; Lichstein, 2007). Pairwise genetic distances between all locations (Weir and Cockerham *F*_ST_, Weir and Cockerham, 1984) were calculated using the R package diveRsity (Keenan et al., 2013), and pairwise shortest geographic distances by sea between all locations, with the R package marmap (Pante and Simon-Bouhet, 2013).

Isolation-by-resistance (IBR) was then investigated, to check for further effects of some seascape features (bathymetry, distance-to-habitat) on the dispersal of grey reef shark populations. IBR models assume a linear relationship between pairwise genetic distance and pairwise resistance distance, a metric both taking into account geographic distance and landscape features between locations (McRae and Beier, 2007).

Furthermore, we used IBR models implementing electrical Circuit Theory (CT) (McRae and Beier, 2007; McRae et al., 2008), and tested their performance in explaining genetic differentiation across our sampling locations of grey reef sharks across the Indo-Pacific. This allowed us to explore different biological hypotheses about gene dispersal for this adult-disperser species. Methods based on CT allow the calculation of a resistance distance between each pair of sampled locations by simultaneously considering all possible pathways connecting these locations, and ascribing resistance values to each pathway.

### 2.5 Resistance maps

Resistance maps were generated in Python (*gdal*) with 10 km cells, based on different hypotheses. The spatial resolution of 10 km was arbitrarily chosen to allow a reasonable computation time, and because grey reef sharks have shown a high residency and relatively small home range of the same order of magnitude (Bonnin et al., 2021; Espinoza et al., 2015a; White et al., 2017). First, we produced a map with homogeneous resistance values across every cell to be used in a ‘CT null model’, corresponding to CT in a homogeneous seascape. Then, bathymetry (GEBCO, gebco.net) and distance-to-habitat resistance maps were drawn separately and in combination as seascape features potentially driving gene flow. We included coral reefs (data.unep-wcmc.org; Spalding et al., 2001) and island nearshores as suitable habitats for grey reef sharks (<u>earthworks.stanford.edu/catalog/harvard-glb-volc</u>). Distance-to-habitat maps were calculated with or without the inclusion of shallow seamounts as suitable habitats (Yesson et al., 2011), selected at a threshold of 280 m corresponding to the reported preferential depth for this species (Last and Stevens, 2009). Different types of relationships between seascape features and resistance were then explored. The grey reef shark being a shallow reef-associated species, we hypothesized that resistance to gene flow was likely to increase with distance-to-habitat and depth, and thus tested multiple values of parameters for linear, logarithmic, and exponential relationships with minimum and maximum thresholds. Different maximum resistance values were also used to calculate the set of resistance maps to run through CT. Full details on resistance maps parametrization are reported in Table S1 and Figure S1.

### 2.6 Pairwise distances using Circuit Theory

The set of obtained resistance maps was then used as input for IBR models using GFLOW (Leonard et al., 2017), an optimized version of the Circuitscape software, estimating pairwise resistance distances between sampling locations using Circuit Theory (McRae, 2006; McRae and Beier, 2007). This batch of obtained pairwise resistance distances between all locations and for every resistance map was then correlated to pairwise genetic distances (pairwise *F*_ST_ estimates linearized using the formula *F*_ST_/(1-*F*_ST_); Figure 2). As a result, in addition to the IBD and CT null models, one IBR model was obtained for each resistance map previously obtained.

**Figure 2.**
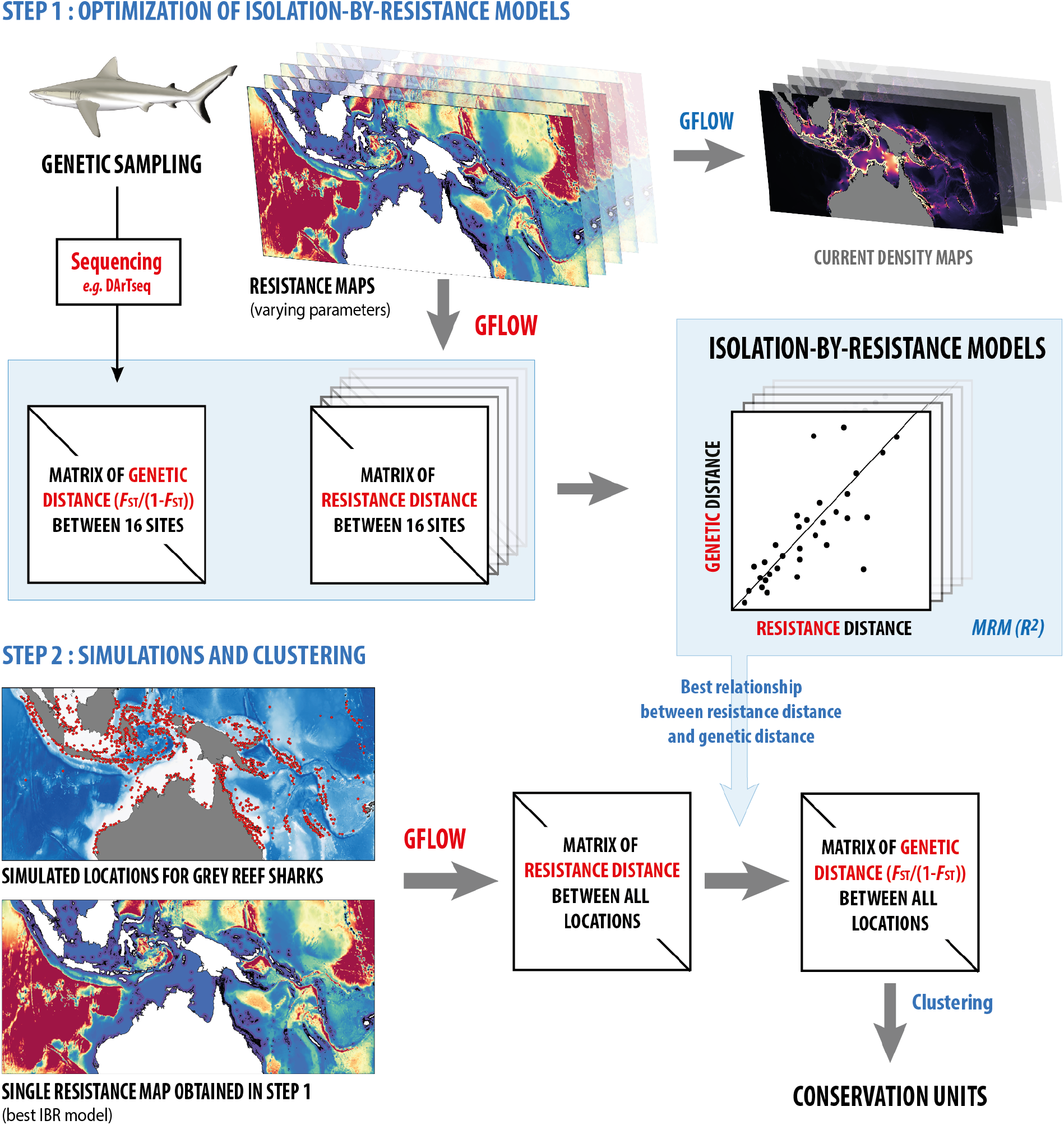
Conceptual framework of the two-step procedure of isolation-by-resistance models to produce conservation units based on simulated locations. In Step 1, numerous resistance maps are produced via the parametrization step described in Figure S1 and Table S1, then used as input in GFLOW, resulting in one matrix of resistance distance per resistance map. For each map corresponding to one parameter set, resistance values are then linked by dyad to genetic distance values (linearized *F*ST), and model robustness evaluated with the regression coefficient (*R*^2^) obtained from a multiple regression on distance matrices (MRM). In Step 2, GFLOW is run on simulated locations for grey reef sharks (Figure S3) with the resistance map corresponding to the best IBR model selected in Step 1. The output, a matrix of resistance distance between all simulated locations, is then converted to genetic distances based on the best IBR linear model. The subsequent matrix of simulated genetic distance is then submitted to a density peaks clustering method described in Rodriguez & Laio (2014).

Models were compared and the best model was chosen by looking at a combination of *R*^2^ values from multiple regression matrices (MRMs), Mantel tests (Mantel, 1967) and AICc (R package AICcmodavg). Because of the high correlation between bathymetry and distance-to-habitat, tests like partial Mantel (Smouse et al., 1986) and MRMs ran independently on the two variables are not optimal (Legendre and Fortin, 2010; Peterman and Pope, 2021). Univariate models were optimized on one hand, representing the best correlations between seascape features independently and genetic distances. On the other hand, multivariate models accounting for both bathymetry and distance-to-habitat cumulated in single resistance maps were also tested with multiple combinations of maximum resistance, relationship shapes and associated parameters. More details on the general framework, parametrization and model optimization are available in Figure 2, Figure S1 and Table S1.

### 2.7 Simulations

Locations with possible presence of grey reef sharks were identified following the same criteria described before as suitable habitat (excluding seamounts) and subsampled at different scales (Froese et al., 2010). The first scale was at the extent of our sampling locations, from the Cocos Keeling Islands to the Eastern New Caledonian volcanic islands. Locations with at least 50 km separation were randomly chosen based on a Matérn process maximizing the number of chosen points (Kiderlen and Hörig, 2013). Next, at the scale of the entire distribution range of the species (tropical Indo-Pacific), including hypothetical presences only based on habitat suitability (ranging between 32°E-130°W longitude and 30°S-30°N latitude), further locations separated by a distance of at least 100 km were randomly chosen using the same method. Such distances between locations were chosen to keep computation time reasonable. The extent to which locations were randomly chosen was narrower than the extent of the resistance maps used as input from GFLOW. Indeed, the artificial boundaries created by the edges of a map can have a non-negligible impact on the calculations of landscape resistance to gene flow (Koen et al., 2010).

The best CT model obtained with our optimization framework, and thus the resistance map best explaining genetic differentiation between the sampling locations, was further used to run GFLOW on the randomly chosen locations (Figure 2). It allowed computation of pairwise resistance distances between all possible locations for grey reef shark presence previously selected with the Matérn process. Such pairwise resistance distances were then converted to pairwise genetic distances (predicted Weir and Cockerham *F*_ST_), using the best relationship between distance matrices obtained in the optimization step (*i*.*e*. from empirical data).

### 2.8 Clustering of subpopulations

The obtained dissimilarity matrices of genetic distances were then subjected to the clustering procedure developed by Rodriguez and Laio (2014) to delineate conservation units (CUs) at both scales. Based on the automatic identification of local density peaks, this method allows the detection of clusters and outliers based on the distance between data points. It is similar to density-based algorithms such as DBSCAN (Ester et al., 1996), however it delineates clusters without introducing a noise-signal cutoff, thus decreasing the probability of low-density clusters being classified as noise (Rodriguez and Laio, 2014). The only variable parameter *d*_c_ was fixed so that the average number of neighbors represents 5% of the total number of points in the dataset (Du et al., 2016; Rodriguez and Laio, 2014). Results produced by this clustering method are very robust across variation of this parameter, particularly on large-scale datasets (Xu et al., 2020).

## 3 RESULTS

OutFLANK and FLK tests identified 8 shared outlier loci, that we excluded before further analyses, leaving a total of 4,983 SNPs considered as neutral.

The discriminant analysis (DAPC) indicated that grey reef shark populations could be split into four distinct clusters (Figure 3A-C), also identified by fastSTRUCTURE (K=4 genetic clusters, Figure 3D). This clustering revealed greater differentiation between areas separated by large distances or deep waters (Figure 3A-C). Sharks sampled at the far remote Chagos showed greater genetic differentiation compared to other sampling locations (Figure 3A). This effect was also observed in populations from the oceanic island of Matthew on the New Hebrides Plate, and the Cocos Keeling Islands, which are both isolated coral reef islands separated by deep oceanic waters (Figure 3B). The DAPC also suggested that sharks from the remote reefs of Chesterfield were more related to sharks from the Great Barrier Reef (GBR) than to sharks from the rest of the New Caledonian archipelago (Figure 3C). Pairwise genetic distances (Weir and Cockerham *F*_ST_ values) confirmed the patterns of differentiation observed by DAPC and fastSTRUCTURE (Table S2).

**Figure 3.**
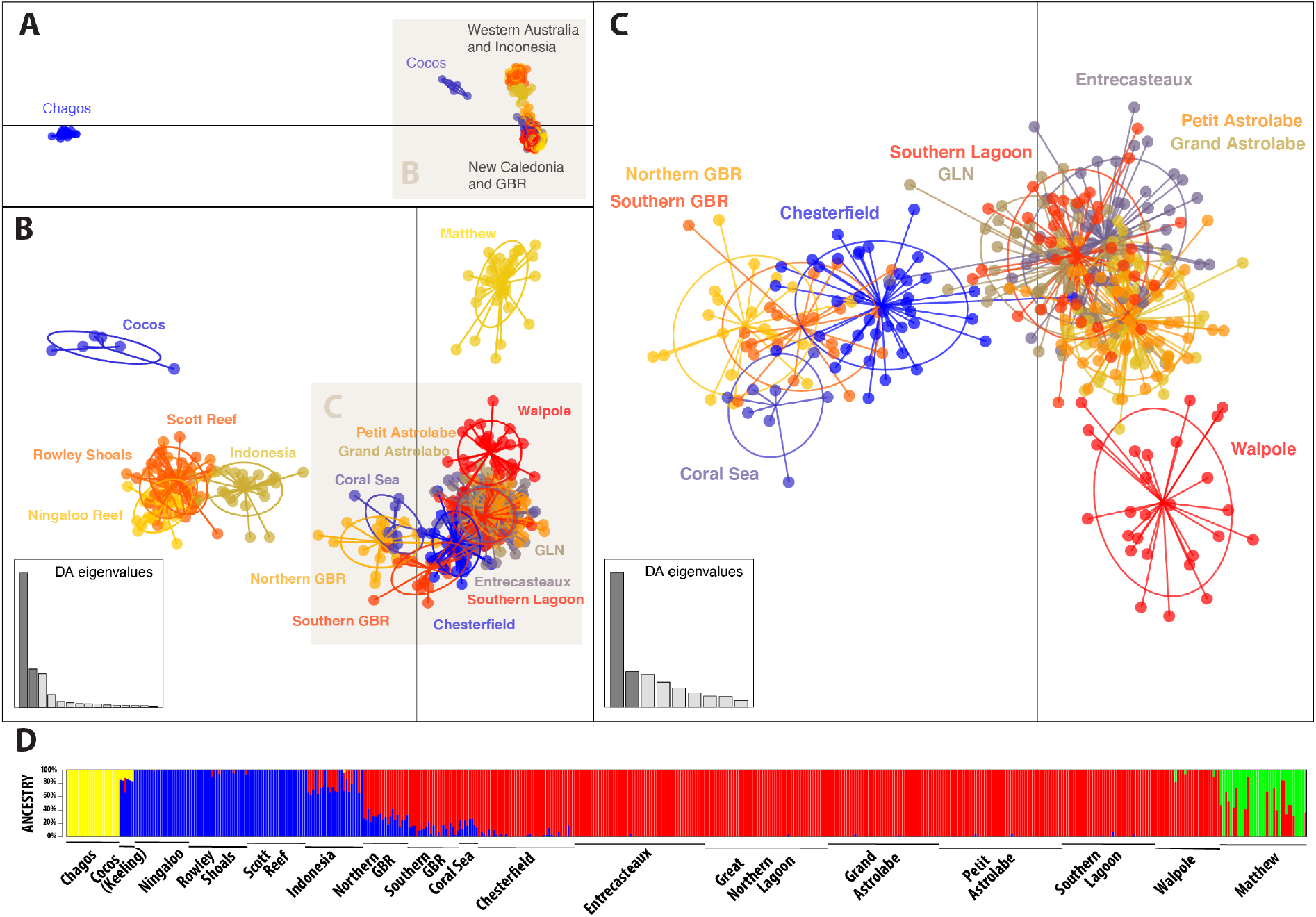
Results from Discriminant Analysis of Principal Components (DAPC) performed on the set of 4,983 filtered neutral SNP data (excluding outliers) from all locations (A), all locations except Chagos (B), and locations in the Pacific Ocean including Eastern Australia and New Caledonia except Matthew (C). Colors and inertia ellipses correspond to sampling locations. (D) Results from fastSTRUCTURE using simple prior and 4 clusters. Samples from Chagos were included in this analysis.

During model optimization, 618 resistance maps were obtained when testing single parameter hypotheses (bathymetry or distance-to-habitat; with or without including seamounts as suitable habitat). Likewise, a total of 93,632 resistance maps combining both seascape features in every possible combination were obtained and used as input for GFLOW.

Isolation-by-distance (IBD) calculated with linear geographic shortest distances between sampling locations explained an important part of genetic differentiation (*R*^*2*^ = 0.493), as well as IBD calculated with Circuit Theory (GFLOW null model, pixels of value 1, *R*^*2*^ = 0.284), but uncertainty and many outliers remained (Figure 4). The distance-to-habitat- and bathymetry-based univariate models, respectively a model with high resistance value when at more than 200 km from any suitable habitat and a model with low resistance at depths shallower than 2000 m, attaining very high resistance values at depths below 4000 m, were highly predictive (*R*^*2*^ = 0.952; Mantel = 0.976; AICc = -904 and *R*^*2*^ = 0.985; Mantel = 0.992; AICc = -1041, respectively). The best model combining both bathymetry and distance-to-habitat was even more predictive (*R*^*2*^ = 0.988; Mantel = 0.994; AICc = -1070). This best model did not include seamounts as suitable habitat and suggests that deep oceanic waters represent a strong barrier to dispersal. It also suggests that habitat proximity of less than 50 km promotes gene flow (Figure S2).

**Figure 4.**
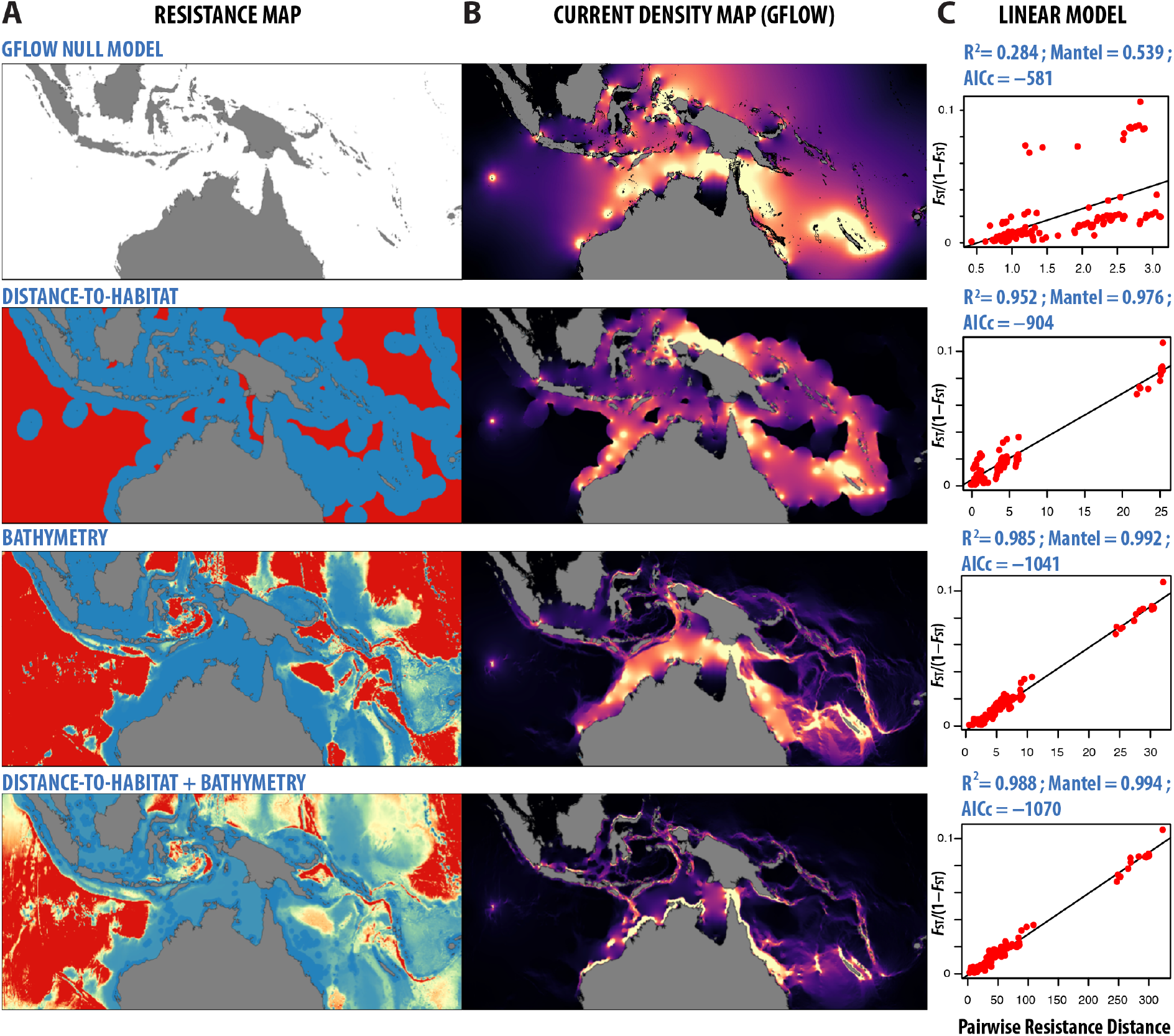
Results of the optimization framework showing (A) the best resistance maps associated with different hypotheses: ‘GFLOW Null Model’, distance-to-habitat alone, bathymetry alone, and the best combined model (distance-to-habitat and bathymetry, without including seamounts in the distance-to-habitat layer); (B) the associated GFLOW maps representing current flowing between every pair of locations for the given resistance map. Brighter colors indicate higher current flow; (C) the subsequent linear relationships between pairwise resistance distances obtained with GFLOW and linearized FST for every pair of locations. *R*^2^, Mantel statistic and AICc are indicated for each model.

At the scale of the entire distribution range, the GFLOW run using the best resistance map produced a matrix of 480,690 pairwise resistance distances between the 981 simulated locations that were randomly selected across the Indo-Pacific, separated by 100 km (Figure S3). Density peaks clustering revealed a total of 38 conservation units comprising ≥ 2 locations, along with 202 isolated locations (Figure 5A). The widest unit was comprised of reefs and oceanic islands in the eastern part of the Indo-Australian plate, along with the southeastern part of the Eurasian plate (Sunda plate), while the western frontier of the unit delineated by the Java Trench. Another wide unit connected reefs from the Solomon and Bismarck plates, while remote islands in the southern part of the Solomon Islands were connected to Vanuatu. Interestingly, Tonga, Fiji, Wallis and Futuna, as well as the southern islands of Tuvalu formed a major unit in the Western Pacific Ocean. Five distinct units encompassed reefs from the Red Sea and the Oman Sea/Persian Gulf, while the western coast of Madagascar and the Comoros, including Mayotte, in the Mozambique Channel were part of a single unit together with a wide section of the eastern coast of Africa. The Seychelles formed a single unit, which was also the case of Chagos. Another unit in the Indian plate was composed of surrounding reefs in India and Sri Lanka, the Laccadive Islands and the Maldives. Lastly, except from some wide units comprised of archipelagos like for instance the western part of Micronesia, or the Tuamotu archipelago in French Polynesia which formed single units, reefs and oceanic islands from the Pacific plate were much fragmented, with the largest proportion of small units and completely isolated patches of habitat (*e*.*g*. Cocos Keeling Islands).

**Figure 5.**
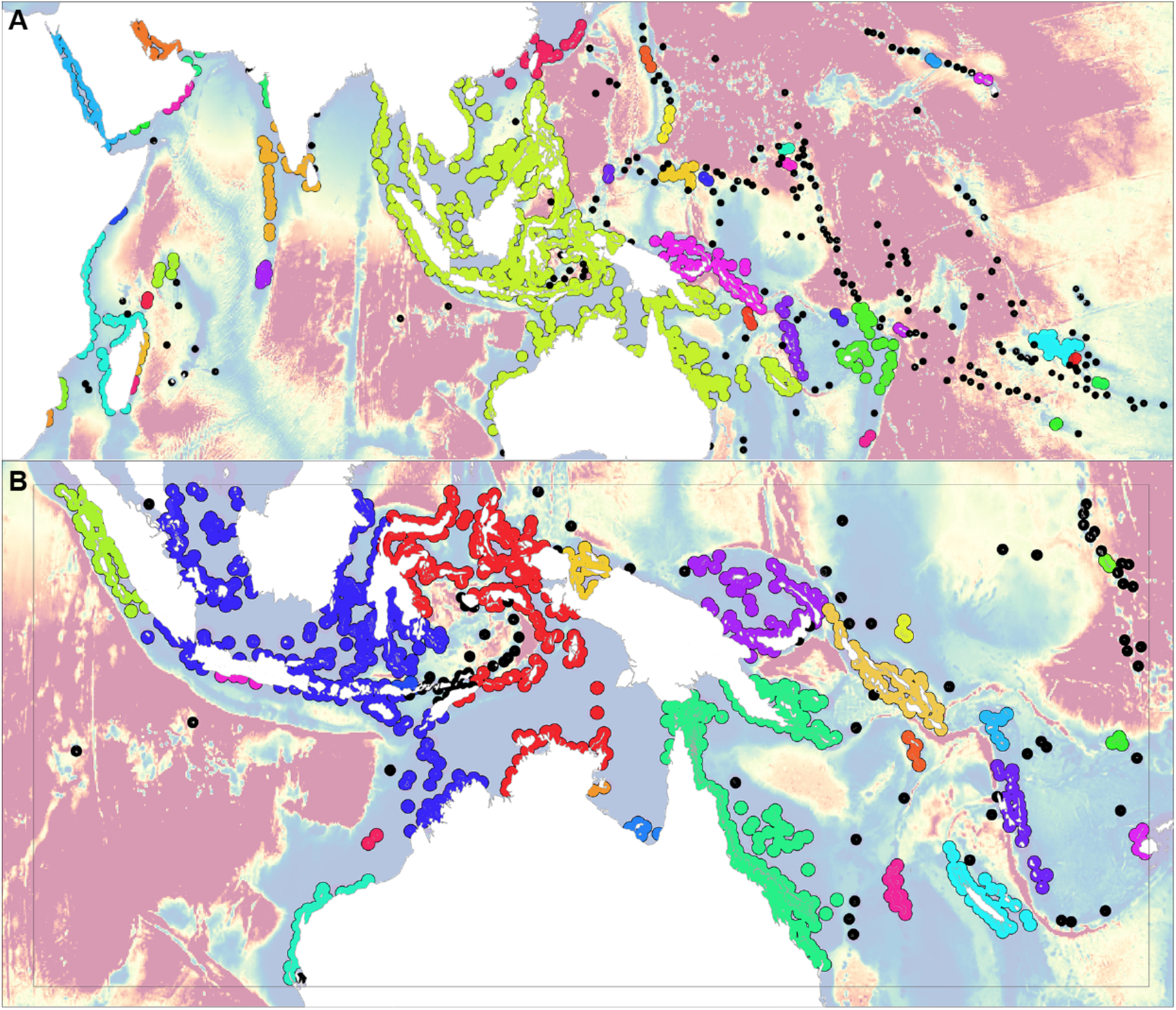
Resistance maps showing the delineation of conservation units (same color), and buffers of radius equal to (A) 100 km at the scale of the whole distribution range of the grey reef shark and (B) 50 km at the scale of our sampling extent. Isolated units comprised of a single location are shown in black with a smaller radius. The grey rectangle on (B) corresponds to the extent on which locations were simulated. Color scale in the background corresponds to resistance values from Figure S2, red corresponding to high values, blue to low values.

At the smaller scale of our sampling extent, GFLOW similarly produced a matrix of 402,753 pairwise resistance distances between the 898 simulated locations separated by 50 km, randomly selected during the Matérn process (Figure S3). Density peaks clustering revealed a total of 21 units (≥ 2 locations) at the scale of our sampling extent, along with 81 isolated locations (Figure 5B). At this scale, the single unit comprising mostly reefs from the Indo-Australian plate was fragmented into several conservation units. Noticeable ones in terms of conservation comprised a western Australian unit, a separate unit constituting the Rowley Shoals, while western Indonesia and northwestern Australia were grouped into a single unit, joined together by Scott Reef and Ashmore and Cartier Islands. Eastern Indonesia and northern Australia off the Northern Territory constituted another very close unit with the east Timor Sea and Arafura Sea acting as corridors. The GBR, along with reefs from the Coral Sea, was connected to Papua New Guinea via the Torres Strait. Reefs from the Bismarck Sea (northern Papua New Guinea) formed a single unit, as well as most Solomon Islands reefs. Interestingly, the Chesterfield Reefs, belonging to New Caledonia, formed a unit by themselves. The rest of the New Caledonian archipelago also formed a single unit, except Matthew and Hunter Islands that were isolated on the far east side of the archipelago, and the Petrie atoll, isolated in the north-east of the main island. Further east, Vanuatu was separated from New Caledonia by the New Hebrides Trench. Among the 81 isolated locations identified by the clustering algorithm, Cocos Keeling and Christmas Islands in the Indian ocean, reefs in the Banda Sea (southeast Asia), Nauru, Tuvalu, Kiribati and other remote islands of the Pacific, as well as remote reefs in the Coral Sea were identified.

## 4 DISCUSSION

Common approaches in landscape or seascape genetics usually focus on genetic connectivity *per se* and propose *ad hoc* explanations based on coincident landscape features (Hirschfeld et al., 2021; McRae and Beier, 2007), therefore hindering the potential of genetic studies to inform conservation planning. Conversely, this study is the first to date linking fine-scale seascape and genetic connectivity of a species with *a priori* testing of hypotheses, followed by predictions at the entire range of a species. Here, with the development of an analytical framework using a custom pipeline that could be applied to a variety of different species and ecosystems, we show its potential for the delineation of hierarchical conservation units at various scales for more targeted protection measures. Applying this methodology for multiple species could provide key information for high resolution management scenarios, particularly for the implementation of MPAs and MPA networks of improved effectiveness (Momigliano et al., 2015). It provides a complementary approach to other modelling frameworks based on movement data from individuals (Martín et al., 2020), and represents an efficient means to predict large-scale conservation units.

Our results reveal that geographic distance is a poor predictor of the genetic structure of grey reef sharks. While genetic and geographic distance are correlated (*R*^2^ = 0.493), the explanatory power of this null model is low compared to IBR models accounting for seascape features (best model *R*^2^ = 0.988). This is not surprising as two underlying assumptions of IBD are clearly unrealistic. The first and most important assumption of IBD is that dispersal occurs through a homogeneous seascape. Grey reef sharks are habitat specialists, being associated almost exclusively with coral reefs (particularly exposed outer slopes) and rocky shoals (Chin et al., 2010; Espinoza et al., 2014; White et al., 2017). Therefore, their dispersal is likely constrained by the availability of suitable habitats (Espinoza et al., 2015b, 2015a). This is further supported by the distribution of clusters along the first two axes of the DAPC (Figure 3), displaying a hierarchical islands structure, typical of a stepping stone model of dispersal (Jombart et al., 2010). Another assumption of IBD and more specifically of Least Cost Path (LCP) is that sharks use direct pathways between locations. This assumption has been shown to bias inference in many organisms (McRae and Beier, 2007), and in the case of grey reef sharks, there is scarce evidence of direct long-distance migration pathways (Bonnin et al., 2019, 2021).

We show that the best model explaining the genetic differentiation of grey reef sharks is one supported by CT and with a very low resistance associated to waters at less than 50 km from optimal habitats, but with very high resistance associated with deep oceanic waters acting as barriers to dispersal. Although the majority of grey reef sharks have been found to be highly resident, some individuals are known to travel large distances across the open ocean (Espinoza et al., 2015a; White et al., 2017). Altogether, our results are congruent with previous studies highlighting that large MPAs (> 50 km) could be effective for a substantive proportion of individuals (Bonnin et al., 2021; Dwyer et al., 2020; Edgar et al., 2014; MacNeil et al., 2020), even though a small number of individuals may disperse further using contiguous habitat patches as travel routes to avoid high resistance barriers such as deep oceanic waters (Bonnin et al., 2019, 2021). We recognize that expanding our modelling wider than the sampling extent (*i*.*e*., the central Pacific) can be problematic, and we call for extensive genetic sampling at a wider scale to confirm these findings. Nevertheless, studies have yet to demonstrate that sharks from unsampled regions have different dispersal behaviors than the Indo-West-Pacific sharks sampled here.

Based on empirical evidence provided by genetic data, our results prove that at a time scale of several generations, a small number of sharks dispersing genes via migration (Bonnin et al., 2021) may have a significant impact on the global genetic structure of the species, and consequently in maintaining standing genetic variation and inbreeding connectivity (sufficient gene flow to avoid harmful effects of local inbreeding, Lowe and Allendorf, 2010) among and between conservation units (CUs). There is, however, an important consideration to be made: *F*_ST_ is a proxy of migration only when populations are at migration-drift equilibrium. Given the long generation time of grey reef sharks (16.4 years, see Robbins, 2006), the young age of some of the sampled habitats (like the GBR), and the evidence of recent population expansions in other coral reef associated requiem sharks (Maisano Delser et al., 2016, 2018), this assumption could be considered as invalid. Assuming all populations have a recent history, as in the closely related *C. melanopterus* (Maisano Delser et al., 2016, 2018), *F*_ST_ is still expected to be correlated to *N*_e_m, but defining CUs using a cut-off based on *F*_ST_ may be misleading: if populations are not at migration-drift equilibrium, *F*_ST_ may be much lower than expected for a given *N*_e_m. A possible solution to such limitation would be to estimate migration rates without assuming equilibrium using the coalescent, or approximations of the coalescent, within an approximate Bayesian computation or composite likelihood approach for parameter estimation (Beaumont et al., 2002; Excoffier et al., 2013, 2021; Gutenkunst et al., 2009; Jouganous et al., 2017). A framework that incorporates IBR models and direct estimates of migration rates based on coalescent simulations would be a significant step forward in seascape genetics, potentially enabling the estimation of much higher migration rates than *F*_ST_-based methods, while taking into account the demographic history of all populations. There are however potential issues to consider. As Momigliano et al. (2021) recently demonstrated, unaccounted demographic events may cause strong biases in parameter estimation, although migration rates are among the least affected demographic parameters.

Protecting threatened mobile species requires a better knowledge of habitat fragmentation and physical barriers in the seascape (Hirschfeld et al., 2021). While the concept of ‘populations’ is used to guide management policy, it covers multiple definitions (Waples and Gaggiotti, 2006) but is often approached by the identification of CUs (Funk et al., 2012). There are also various definitions of CUs in the scientific literature, with a major distinction between Evolutionary Significant Units (ESUs) and Management Units (MUs), but there is a consensus on the fact that identifying CUs is a crucial first step for the conservation of wild populations (Barbosa et al., 2018; Funk et al., 2012). CUs are also recognized as being hierarchical, with units at wider scale comprising multiple smaller units (Barbosa et al., 2018; Funk et al., 2012; Weckworth et al., 2018). The investigation of local genetic and demographic additional clues (*i*.*e*. genetic diversity, *N*_e_, relative abundance) might help to better delineate units and take more appropriate management measures (Barbosa et al., 2018; Domingues et al., 2017).

One of the important aspects of our results is that the defined CUs, even at local scale, generally encompass the Exclusive Economic Zones (EEZs) of multiple countries. As such, conservative spatial planning would require coordinated international efforts (Harrison et al., 2018; Mackelworth et al., 2019). Moreover, slowing the ongoing decline of natural populations of mobile species like sharks requires not only scientific collaborations, but also support from managers and policy-makers across borders (Dunn et al., 2019; Sequeira et al., 2019). A further issue impacting mobile predators such as the grey reef shark is the fragmentation of populations observed through a high proportion of putative conservation units represented only by a single location of suitable habitat. This highly fragmented pattern holds true for the two hierarchical scales (81 of 103 units at the scale of our sampling extent, 202 of 240 at the entire distribution range of the species), knowing that the clustering algorithm used is conservative in the number of detected outliers (Rodriguez and Laio, 2014). Special attention should thus be given by managers to such isolated locations that deserve high conservation priority, hosting populations potentially vulnerable to anthropogenic pressures such as harvesting, with a low capacity of rebuilding populations via migration and subject to inbreeding depression for depleted populations (Kardos et al., 2018; Ralls et al., 2018).

## 5 CONCLUSIONS

We developed and used a predictive modelling framework to infer barriers to gene flow and map the connectivity of grey reef sharks across the Indo-Pacific. We provide novel insight on the conservation of this marine predator by estimating connectivity beyond sampled locations and by delineating hierarchical conservation units. We conclude that the distribution and movement of grey reef sharks are reliant on more than just geographical availability and demonstrate the importance of using this framework for the integration of genetic connectivity in the field of marine spatial planning. Our findings are not limited to grey reef sharks, and this framework can be applied to any adult disperser species. Hence, we call for the use of this approach to better understand dispersal patterns of other marine species at different scales. We recommend including such information alongside ecological data, habitat use, and governance of areas used when considering management strategies, and even applying this framework on multiple species as part of a systematic and integrated conservation planning approach (Sala et al., 2021).

## Acknowledgments

We thank the crew of the Research Vessel Amborella (Government of New Caledonia), the many volunteers that contributed to field trips in New Caledonia, and Laurent Millet of the genetic lab at IRD Nouméa for help in the processing of samples. We appreciate the support of Pierre-Edouard Guerin (CEFE/CNRS) for SNPs filtering and of Jérôme Lefèvre (ENTROPIE) and Eric Mermet (EHESS) on map computation.

## Notes

**Conflict of interest** The authors declare no conflict of interest.

**Funding** Sampling in New Caledonia was funded by research grants of Laurent Vigliola (Programme APEX, Government of New Caledonia, Pew Charitable Trust, Total Foundation, IRD core funding), and David Mouillot (Programme APEX, Total Foundation). Data collection in other countries was supported by the Great Barrier Reef Marine Park Authority, Cooperative Research Centre (CRC) Reef, Australian Coral Reef Society, and PADI Aware. This research was funded also by the Sea World Research and Rescue Foundation and the Academy of Finland (grant 316294 to PM).

### Competing Interest Statement

The authors have declared no competing interest.

