## Supporting Information for "Identifying barriers to gene flow and hierarchical conservation units from seascape genomics: a modelling framework applied to a marine predator"

**Table S1.** Formulas and parameter values used for exponential, logarithmic, and linear relationships between bathymetry and/or distance-to-habitat and resistance to gene flow. See Figure S1 for a graphical example showing how the two variables are combined to generate resistance maps. $R_{max}$ is used as a simple multiplier in univariate maps and combinations of $R_{max}$ values are tested (Figure S1) in bivariate maps.

| | **BATHYMETRY** | **DISTANCE-TO-HABITAT** |
|---|---|---|
| **EXPONENTIAL** | y = exp(a*(x/c-1)) for x ≤ c ; y = 1 for x ≥ c | |
| c | 100, 1000, 3000, 4000, 5000 | 100, 250, 500, 2000 |
| a | 2, 6, 8, 10, 32 | 2, 8, 32 |
| $R_{max}$ | 10, 50, 75, 100, 125, 750, 1000, 1250, 10000 | 1, 5, 10, 15, 50, 75, 100, 1000 |
| **LOGARITHMIC** | $y = \log_{10}(a*x+1) / \log_{10}(c*a*x+1)$ | |
| c | 8000 | 3000 |
| a | 0.001, 0.01, 0.1, 1 | 0.001, 0.01, 0.1, 1 |
| $R_{max}$ | 10, 50, 100, 1000, 10000 | 1, 5, 10, 15, 50, 75, 100, 1000 |
| **LINEAR** | y = 0 for x ≤ $C_{min}$ ; y = 1 for x ≥ $C_{max}$ ; y = x / $C_{max}$ for $C_{min}$ ≤ x ≤ $C_{max}$ | |
| $C_{min}$ | 0, 300, 1000 | 0, 300, 1000 |
| $C_{max}$ | 1000, 4000, 8000 | 1000, 4000, 8000 |
| $R_{max}$ | 10, 50, 100, 1000, 10000 | 1, 10, 50, 100, 1000 |

**Table S2.** Pairwise measures of genetic differentiation (Weir and Cockerham $F_{ST}$) between all the sampling locations except Chagos.

| | Cocos | Ningaloo Reef | Rowley Shoals | Scott Reef | Indonesia | Northern GBR | Southern GBR | Coral Sea | Chesterfield Reefs | Entrecasteaux | Great Northern Lagoon (GLN) | Grand Astrolabe | Petit Astrolabe | South NC | Walpole |
|---|---|---|---|---|---|---|---|---|---|---|---|---|---|---|---|
| Ningaloo Reef | 0.067 | | | | | | | | | | | | | | |
| Rowley Shoals | 0.068 | 0.0021 | | | | | | | | | | | | | |
| Scott Reef | 0.064 | 0.0025 | 0.0016 | | | | | | | | | | | | |
| Indonesia | 0.068 | 0.0055 | 0.0054 | 0.0039 | | | | | | | | | | | |
| Northern GBR | 0.073 | 0.0143 | 0.0141 | 0.0125 | 0.0080 | | | | | | | | | | |
| Southern GBR | 0.079 | 0.0200 | 0.0183 | 0.0175 | 0.0126 | 0.0048 | | | | | | | | | |
| Coral Sea | 0.077 | 0.0169 | 0.0159 | 0.0144 | 0.0097 | 0.0031 | 0.0032 | | | | | | | | |
| Chesterfield Reefs | 0.081 | 0.0193 | 0.0189 | 0.0175 | 0.0128 | 0.0064 | 0.0039 | 0.0035 | | | | | | | |
| Entrecasteaux | 0.081 | 0.0215 | 0.0204 | 0.0196 | 0.0135 | 0.0082 | 0.0078 | 0.0085 | 0.0048 | | | | | | |
| GLN | 0.080 | 0.0205 | 0.0202 | 0.0189 | 0.0131 | 0.0070 | 0.0074 | 0.0080 | 0.0043 | 0.0010 | | | | | |
| Grand Astrolabe | 0.081 | 0.0215 | 0.0210 | 0.0197 | 0.0138 | 0.0088 | 0.0090 | 0.0090 | 0.0066 | 0.0026 | 0.0020 | | | | |
| Petit Astrolabe | 0.082 | 0.0216 | 0.0212 | 0.0199 | 0.0144 | 0.0088 | 0.0080 | 0.0081 | 0.0057 | 0.0015 | 0.0015 | 0.0010 | | | |
| South NC | 0.080 | 0.0200 | 0.0203 | 0.0189 | 0.0138 | 0.0080 | 0.0077 | 0.0077 | 0.0051 | 0.0019 | 0.0012 | 0.0025 | 0.0018 | | |
| Walpole | 0.082 | 0.0229 | 0.0214 | 0.0207 | 0.0152 | 0.0115 | 0.0118 | 0.0098 | 0.0084 | 0.0051 | 0.0046 | 0.0049 | 0.0039 | 0.0052 | |
| Matthew | 0.097 | 0.0355 | 0.0341 | 0.0317 | 0.0265 | 0.0223 | 0.0233 | 0.0213 | 0.0195 | 0.0158 | 0.0153 | 0.0155 | 0.0145 | 0.0160 | 0.0128 |

**Figure S1.** Conceptual framework of the parametrization steps used to generate resistance surfaces. (A) Mathematical functions used to model the relationship between depth and/or distance-to-habitat and resistance to gene flow. Exponential, logarithmic, and linear relationships were used, with different parameter values (see Table S1). Bathymetry is shown as an example, but the same relationships were used with distance-to-habitat. (B) Framework showing the combination of depth and habitat with one example of an exponential relationship between bathymetry and resistance, with a ceiling at 4000 meters, a logarithmic relationship between distance-to-habitat and resistance, and a maximum resistance value associated with bathymetry superior to the one associated with distance-to-habitat ($R_{maxB}$ > $R_{maxD}$).

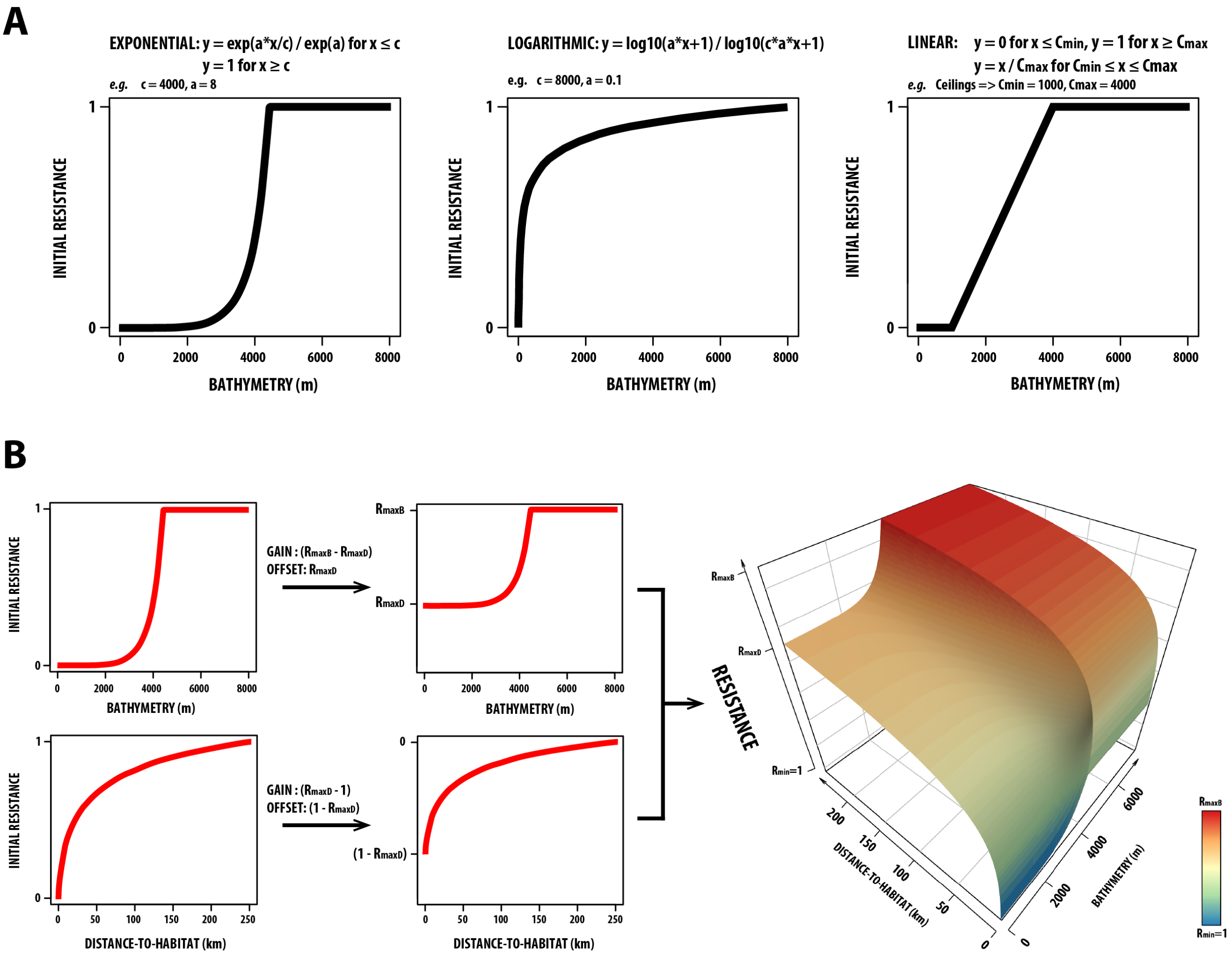

**Figure S2. (A)** Resistance map selected from the best IBR model combining bathymetry and distance-to-habitat, at the scale of the sampling extent and **(B)** relationship between biogeographical features and resistance. This map and the associated relationship were selected after model optimization between resistance values (z-axis) and a combination of bathymetry and distance-to-habitat (x- and y-axes). Minimum and maximum resistance values are 1 and 1000. Resistance attains 50 at locations at more than 50 km from any suitable habitat, and quickly attains high values in deep waters.

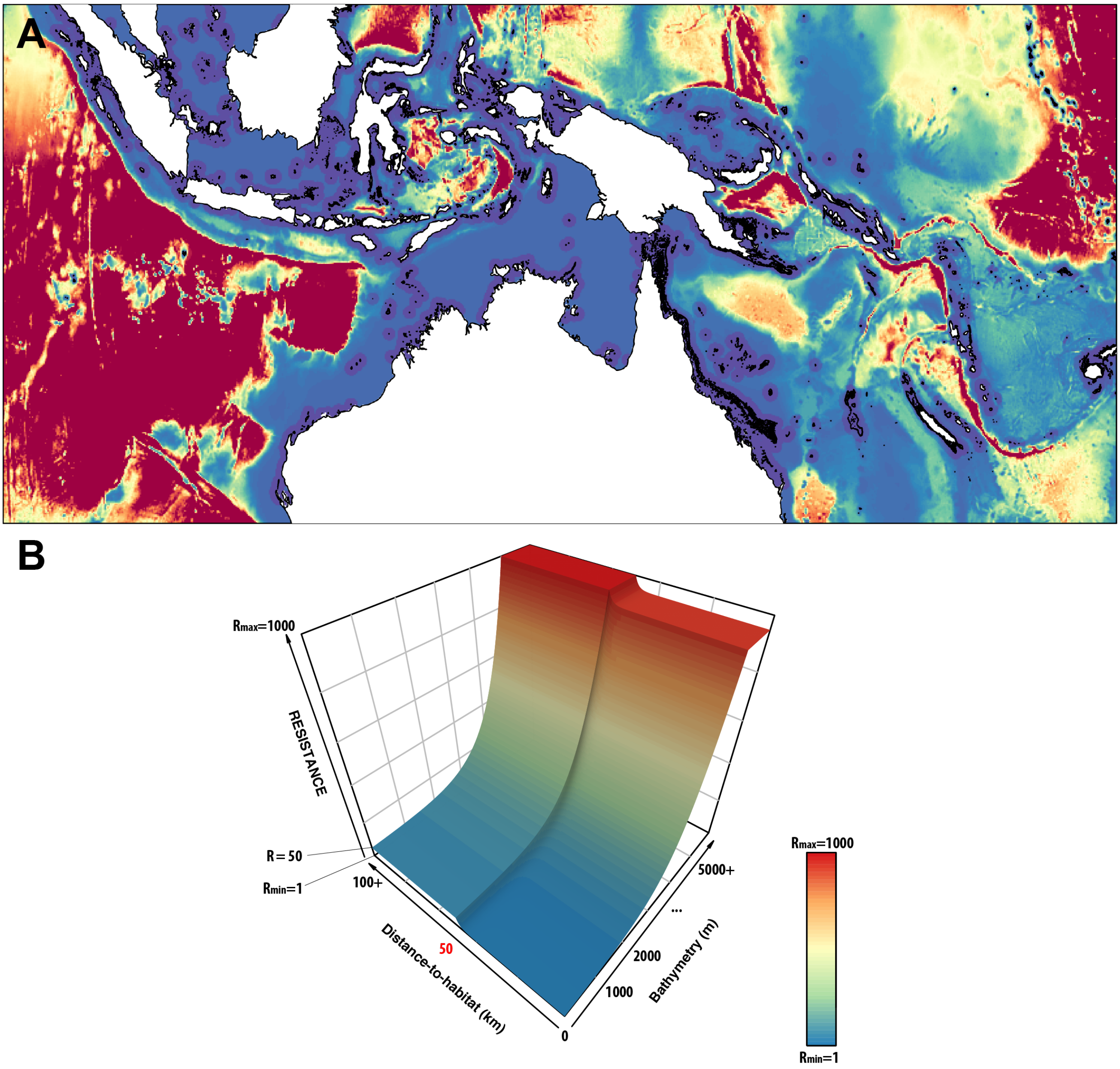

**Figure S3.** Maps of simulated locations randomly chosen via a Matérn process maximizing the number of chosen points. **(A)** At the scale of the sampling extent, 898 locations were selected, separated by a distance of at least 50 km. **(B)** At the scale of the distribution range of the species, 981 locations were selected, separated by a distance of at least 100 km. The yellow extent in (B) corresponds to the scale of the sampling extent in (A).

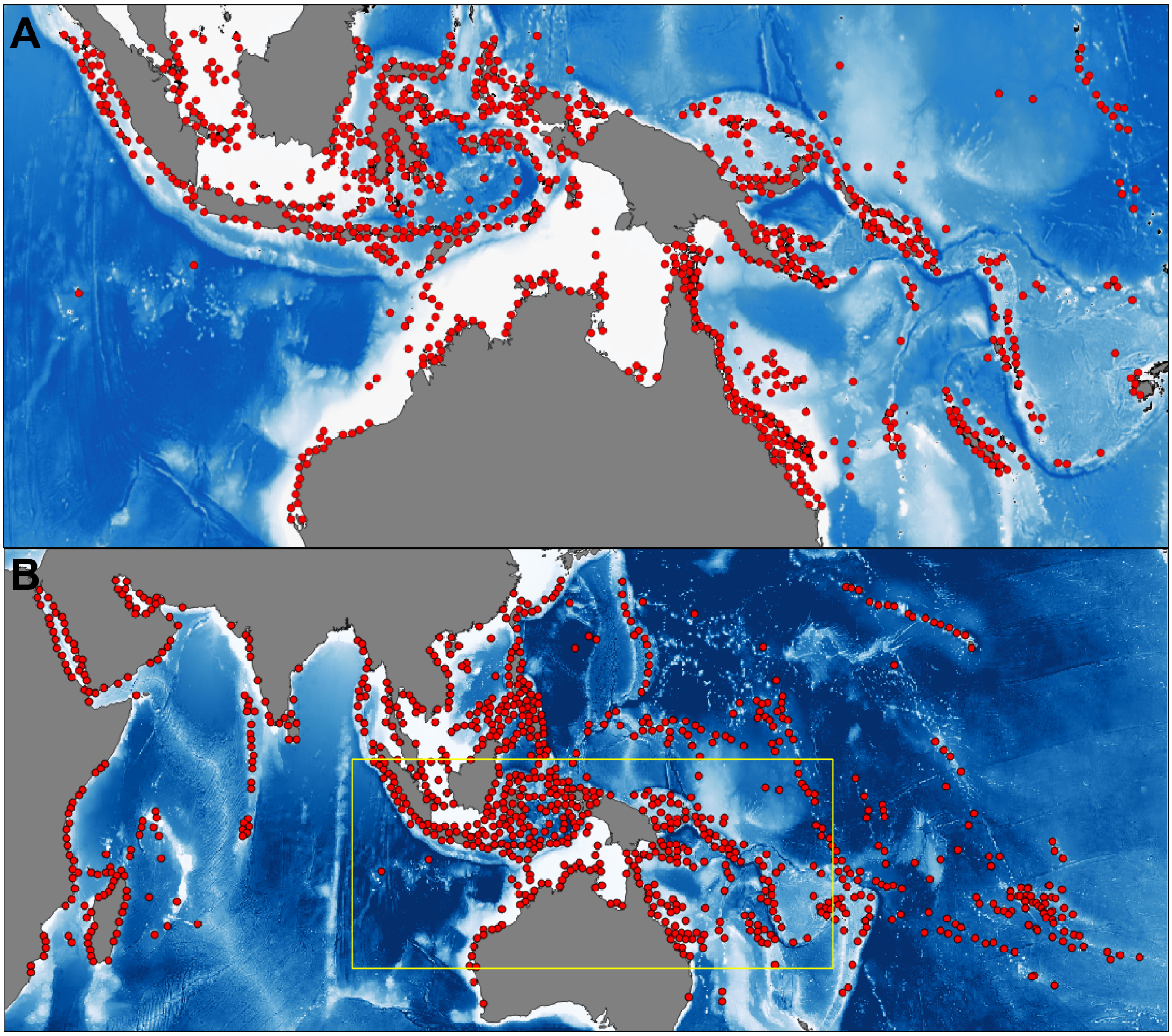
